# Ropinirole, a dopamine agonist with high D_3_ affinity, reduces proactive inhibition: a double-blind, placebo-controlled study in healthy adults

**DOI:** 10.1101/2020.04.27.063560

**Authors:** Vishal Rawji, Lorenzo Rocchi, Tom Foltynie, John C. Rothwell, Marjan Jahanshahi

## Abstract

Response inhibition describes the cognitive processes mediating the suppression of unwanted actions. A network involving the basal ganglia mediates two forms of response inhibition: reactive and proactive inhibition. Reactive inhibition serves to abruptly stop motor activity, whereas proactive inhibition is goal-orientated and results in slowing of motor activity in anticipation of stopping. Due to its impairment in several psychiatric disorders, the neurochemistry of response inhibition has become of recent interest. Dopamine has been posed as a candidate mediator of response inhibition due to its role in functioning of the basal ganglia and the observation that patients with Parkinson’s disease on dopamine agonists develop impulse control disorders. Although the effects of dopamine on reactive inhibition have been studied, substantial literature on the role of dopamine on proactive inhibition is lacking. To fill this gap, we devised a double-blind, placebo-controlled study of 1 mg ropinirole (a dopamine agonist) on response inhibition in healthy volunteers. We found that whilst reactive inhibition was unchanged, proactive inhibition was impaired when participants were on ropinirole relative to when on placebo. To investigate how ropinirole mediated this effect on proactive inhibition, we used hierarchical drift-diffusion modelling. We found that ropinirole impaired the ability to raise the decision threshold when proactive inhibition was called upon. Our results provide novel evidence that an acute dose of ropinirole selectively reduces proactive inhibition in healthy participants. These results may help explain how ropinirole induces impulse control disorders in susceptible patients with Parkinson’s disease.

## Introduction

Response inhibition describes the cognitive processes involved in the suppression of unwanted thoughts and action [1]. It can be divided into reactive and proactive inhibition, whereby reactive inhibition is called upon in response to sudden sensory cues and serves to abruptly stop motor activity, whereas proactive inhibition is goal-orientated and results in slowing of motor activity in anticipation of stopping. A breakdown in response inhibition is a feature of several psychiatric disorders such as schizophrenia [2], addiction [3] and attention deficit hyperactivity disorder [4]. This has encouraged a drive to further understand the neurochemical substrates of response inhibition. Use of dopamine agonist medication in Parkinson’s disease (PD) has been shown to predispose to impulse control disorders (ICDs), manifesting as pathological gambling, hypersexuality and excessive shopping [5]. This observation has therefore prompted investigation of dopamine as a mediator of response inhibition.

A network involving the basal ganglia, right inferior frontal gyrus and pre-supplementary motor area has been implicated in volitional and habitual inhibition [1], and in mediating stopping on behavioural tasks employed to probe reactive and proactive inhibition [6]. Since dopamine is a key neurotransmitter modulating activity in these regions, it seems plausible to study the role of dopamine in response inhibition [7,8]. Infusions of a selective D_1_ receptor antagonist into the rat striatum improves reactive motor inhibition, whereas infusions of a D_2_ receptor antagonist impairs it [9]. In humans, positron emission tomography of striatal D_2_ and D_3_ receptor availability is negatively correlated with the speed of response inhibition and positively correlated with inhibition related functional magnetic resonance imaging activation in the fronto-striatal stopping network [10]. Administration of the dopamine and noradrenaline reuptake inhibitor, amphetamine, increases D_2_ receptor expression and improves measures of reactive inhibition [11].

In addition to the link between D_1_/D_2_ receptors, response inhibition and impulsivity [12], an association between dopamine agonists with D_3_ receptor affinity and ICD generation in PD has been found; with pramipexole and ropinirole being two agents both with relatively high D_3_ affinity compared to other dopamine agonists and greatest risk of ICD generation [13,14]. The link between use of dopamine agonists with relatively high D_3_ affinity and response inhibition remains limited and only reactive inhibition has been explored [15,16]. The effect of these agents on proactive inhibition, on the other hand, has not been investigated.

Our aim was to investigate the effect of dopamine agonists on reactive and proactive response inhibition in healthy participants. We chose the dopamine agonist, ropinirole, which is commonly used clinically and has been implicated in the generation of ICDs in PD. Ropinirole directly activates dopamine receptors with a relatively high affinity for D_3_ receptors [17,18] and hence is a suitable candidate to explore the effect of dopamine agonists on motor inhibition. We devised a randomised, double-blind, placebo-controlled trial to investigate reactive and proactive inhibition in a conditional stop signal task (CSST) under the influence of a placebo or ropinirole.

## Methods

### Participants

Participants were recruited from the staff and students of University College London. Participants were screened to exclude any psychiatric or neurological illness, drug or alcohol abuse and also for any contraindications to ropinirole. A power calculation showed that 30 participants would be needed to show a 20% reduction of the response delay effect (RDE), measure of proactive inhibition, under ropinirole, assuming an intraclass correlation coefficient of 0.655 with 80% power at an alpha level of 0.05. Therefore, 30 healthy volunteers (20 male) aged 19-30 (mean age 23.63, SD 3.64) participated in this study.

### Conditional Stop-signal task (CSST)

The conditional stop-signal task (CSST) is a validated task to probe reactive and proactive inhibition [19–22]. Participants performed four blocks of the CSST driven by a custom-made MATLAB (MathWorks) script using Psychtoolbox. Each block consisted of 102 go trials (51 critical, 51 non-critical) and 34 stop trials (17 critical and 17 non-critical). A trial on the CSST begins with a white fixation cross, which is replaced 500 ms later by one of two imperative stimuli (right or left arrow). The presentation of these arrows is random, and each occurs with 50% probability. The participant is asked to respond as quickly as possible to the right or left imperative stimulus by pressing the ‘M’ or ‘Z’ key on the keyboard with their right or left index fingers, respectively. On 25% of the trials, a stop signal is presented after the go signal, which instructs participants to abort their ongoing movement (stop trial). The timing of this delay between go and stop signals is called the stop signal delay (SSD). The SSD can occur at one of four time points (100, 150, 200 and 250 ms) and was adjusted using a staircase tracking procedure. The SSD for a particular trial was altered based on the outcome of the previous trial; successfully stopping in the previous stop trial would increase SSD by 50 ms (to make the next trial harder to inhibit). Failure to stop on a stop trial would decrease SSD by 50 ms in the next trial (thereby making stopping easier on the next trial). SSD was set to 150 ms at the beginning of each block. Catch trials, where no signals were given, were also presented. At the beginning of each block, participants were told that they would have to follow the stopping rule if the stop signal was presented for one direction (critical direction) but to ignore the stop signal if it appeared after the other imperative signal (non-critical direction). Hence, responses between the critical and non-critical responses could be compared to determine proactive control. The structure of the block was pseudorandomised, such that one stop trial appears in every four trials.

Reaction times were measured as the time interval between the imperative stimulus (right arrow) onset and the button press. Trials were first organised into whether they were to the critical or non-critical direction. For ease, the remainder of this section will refer to ‘right’ as the critical direction and left as the non-critical direction. The critical go reaction time was the reaction time measured on critical (right arrow) go trials and the non-critical go reaction time was the reaction time measured during non-critical (left arrow) go trials. p(inhibit) was calculated as the proportion of successful stop trials (where the participant correctly aborted their response) to the critical (right) direction. The reaction time on failed stop trials (stop trials where the participant failed to stop and hence pressed a button) was also calculated (Stop respond reaction time). The stop-signal reaction time (SSRT) was calculated using the integration method [23,24]. The number of omitted trials were also recorded – omitted trials were ones where no button press was made. Participants slow down their responses in anticipation of potential stopping – which reflects proactive inhibition. To measure this behaviourally, we calculated the response delay effect (RDE) by subtracting the non-critical (left) go reaction time from the critical (right) go reaction time. Conflict-induced slowing (CIS), a measure of temporary braking to stop signals on non-critical trials, was measured by subtracting the non-critical go reaction time (left go trial) from the non-critical stop reaction time (left stop trial). Importantly, participants were told to prioritise responding to go stimuli and not to slow-down in anticipation of a stop-signal.

### Protocol

The study was approved by the Local Ethics Committee (UCL Ethics ID: 9669/002). Written, informed consent was obtained from each participant. After performing one block of the CSST, as a practice block, participants were then given one of two pills (ropinirole 1mg or placebo). Both the participants and the experimenter were blinded and only a third-party investigator, not part of the study, knew the identity of the pills. For each participant, the pill given was selected using a random number generator. The participant stayed in the room for one hour, a time period consistent for ropinirole to reach an appropriate blood level to have CNS effects [25,26]. Following this, the participants performed four blocks of the CSST: two blocks with right as the critical direction and two blocks with left at the critical direction. The order of these blocks was also randomised. After at least 48 hours, the participant returned to the laboratories and underwent the same protocol as session one, except with the other pill.

### Drift-diffusion modelling

A Bayesian hierarchical drift-diffusion model (HDDM) [27] was used to investigate the strategic effects on task performance on the go trials of the CSST when stopping may (critical) or may not (non-critical) be required. DDMs are used to model two-choice, decision-making tasks and aim to resolve the parameters modulated when decisions are made under different circumstances. The model outline is such that activity starts at a starting point (*z)*. After a delay for sensory processing of the imperative stimulus (*n*_*1*_), activity increases towards one of two decision boundaries (*a,* one for each choice) at a rate (*v*). When activity reaches one of the thresholds, the choice is selected; after another delay for motor execution (*n*_*2*_), the choice is made. Usually, the delays for sensory processing and motor execution are combined into one non-decision time (*n*).

We used the HDDM to investigate how ropinirole modulated decision-making parameters during the CSST. Crucially, the HDDM results in better model fits by taking advantage of both the similarity and differences between the participants’ performance; individual participant parameters are drawn from a group posterior such that if participants are similar, the variance in the group distribution is small. We used the HDDM toolbox [27] to estimate DDM parameters. We computed seven different models by varying drift rate, boundary separation and non-decision time between context (critical vs non-critical) and drug (ropinirole vs placebo). The starting point was set to half of the boundary separation since the left/right go cues could appear with equal probability. The optimal model was chosen by using the deviance information criterion (DIC), where a lower DIC indicates a higher likelihood for that model. To maximise available data for the model, we combined all four blocks per drug condition by labelling trials as critical and non-critical, independent of whether they were right or left-handed responses. Markov chain Monte Carlo sampling methods were used to construct the posteriors distribution for parameters (10,000 samples generated, 2000 burn in). To confirm model convergence, we calculated the R-hat (Gelman-Rubin) statistic.

### Data analyses

We were particularly interested in how behavioural inhibition was altered under the influence of the dopamine agonist, ropinirole. To this end, we performed paired t-tests between conditions (ropinirole and placebo) for the following parameters: RDE (proactive inhibition), SSRT (reactive inhibition), CIS, and go discrimination errors. We considered p values of < 0.05 as statistically significant.

Using the HDDM allowed for Bayesian analysis of posterior probability distributions for each parameter, between drug (ropinirole vs placebo) and trial type (critical vs non-critical) conditions. Posterior probabilities of greater than 95% were considered significant.

## Results

### Behavioural measures

Behavioural measurements are shown in Table 1. As expected from the race model, reaction times on failed stop trials (Stop respond) were faster than reaction times on critical go trials for both placebo and ropinirole conditions (placebo: t = 9.20, p < 0.001, d = 0.74, 95% CI [0.21 – 1.25]; ropinirole: t = 3.78, p = 0.001, d = 0.39, 95% CI [−0.13 – 0.89]). There was also an expected go reaction time difference between critical and non-critical trials measured as RDE due to the anticipation of stopping on critical trials (placebo: t = 11.00, p < 0.001, d = 0.99, 95% CI [0.44 – 1.51]; ropinirole: t = 3.78, p = 0.001, d = 0.87, 95% CI [0.33 – 1.39]).

**Table 1:**
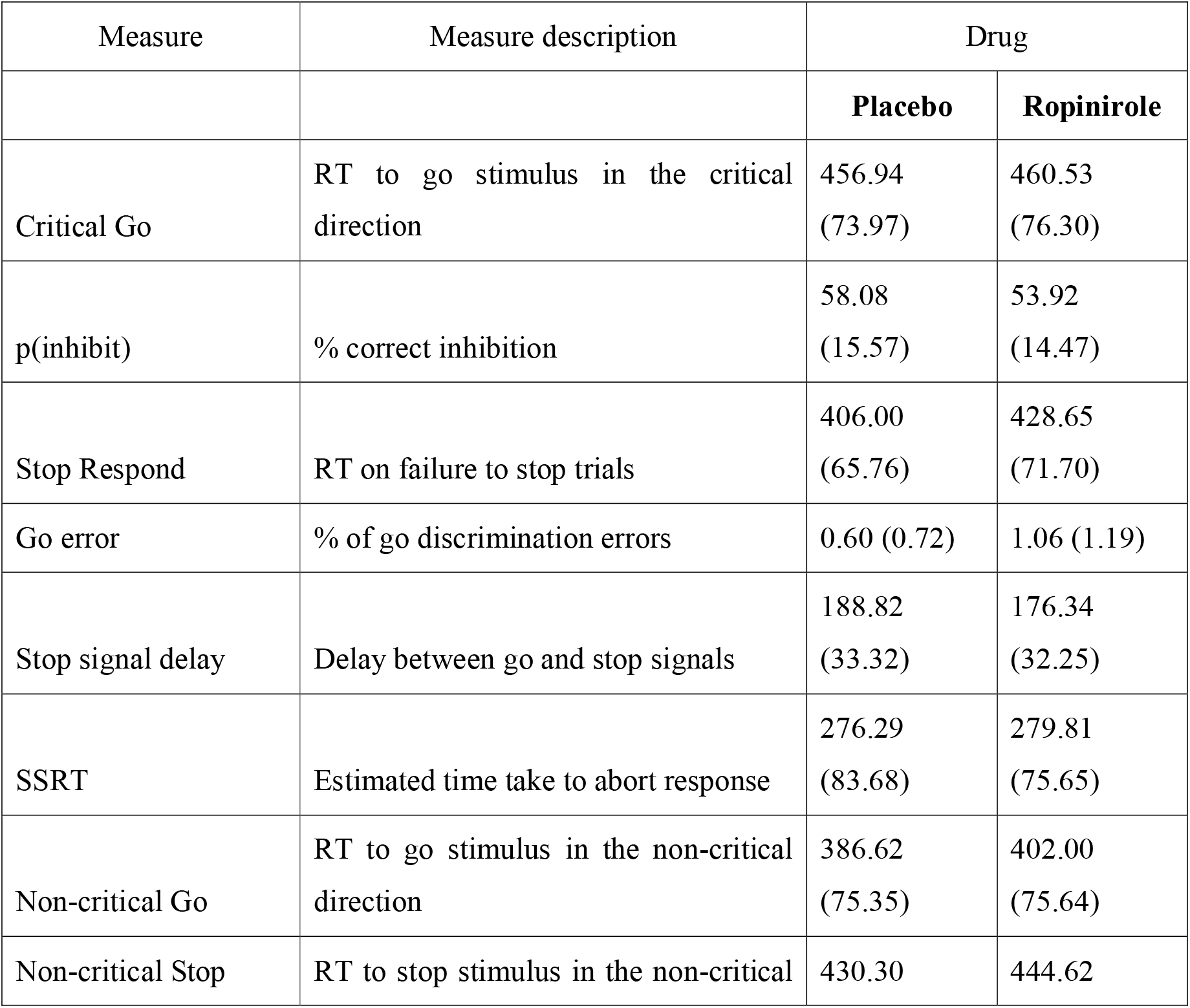
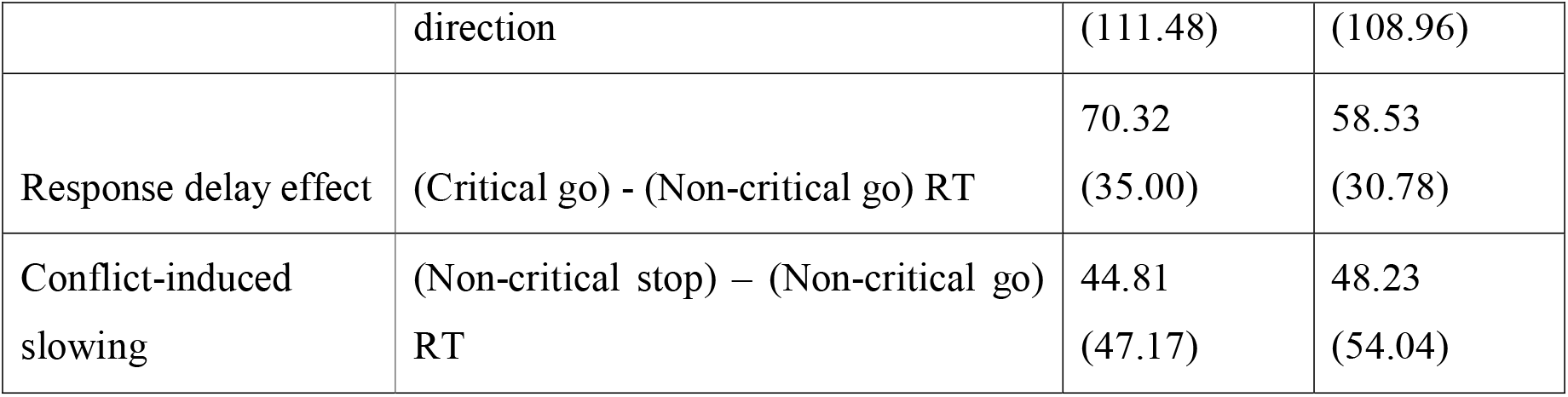
Behavioural measurements from the Conditional Stop-signal task performed by participants on placebo and ropinirole. Mean values are accompanied by SD in brackets. Reaction times are given in milliseconds.

### Ropinirole reduces proactive inhibition and induces more commission errors

We performed a paired t-test between the RDE for placebo and ropinirole and found significantly reduced proactive inhibition on ropinirole (t = 2.25, p = 0.032, d = 0.36, 95% CI [−0.16 – 0.86]). The SSRT is a measure of reactive inhibition; it is the estimated time taken for stopping on presentation of the stop signal. A paired t-test comparing the SSRTs between drug conditions showed that reactive inhibition was unchanged on ropinirole relative to placebo (t = 0.29, p = 0.770, d = 0.04, 95% CI [−0.55 – 0.46]). We also found that under the influence of ropinirole, participants made more go discrimination errors than when on placebo (t = 2.29, p = 0.029, d = 0.47, 95% CI [−0.05 – 0.98]). CIS was unchanged on ropinirole compared to placebo (t = −0.41, p = 0.682, d = −0.07, 95% CI [−0.57 – 0.44]).

### On ropinirole, reduced proactive inhibition is due to failure to raise the decision threshold in anticipation of stop trials

Of the seven models we tested, the one with the lowest DIC was a model allowing boundary separation, drift rate and non-decision time to vary (Table 2); parameter estimation therefore comes from this model. To validate the model, we simulated data for each trial type, under each drug condition, using the parameters from the model with the lowest DIC. The model recapitulated both the mean reaction times (critical go placebo: 461.67 ms, non-critical go placebo: 385.48 ms, critical go ropinirole: 468.41 ms, and non-critical go ropinirole: 400.57 ms) and reaction time distributions for each drug/trial permutation, shown in Figure 2. Figure 3 shows the posterior parameter estimation for the critical and non-critical Go trials in the ropinirole and placebo conditions. We first looked at the effect of trial type: during critical Go trials, when participants were slowing their responses and hence employing proactive inhibition, boundary separation was greater (posterior probability 100%), drift rate was lower (posterior probability 100%) and non-decision time was shorter (posterior probability 99.96%), compared to non-critical Go trials. Turning our attention to the effect of ropinirole, we found that participants’ boundary separation was smaller (posterior probability 100%), drift rate was lower (posterior probability 100%) and non-decision time was smaller (posterior probability 98.66%), compared to when assessed on placebo.

**Table 2:** Deviance information criterion values for each model. The Table shows the DIC value for each model tested. A lower DIC value indicates a greater likelihood for that model. a = boundary separation, v = drift rate, n = non-decision time.

| Model | DIC |
| --- | --- |
| a | -46135.48 |
| v | -46842.23 |
| t | -45503.35 |
| a,v | -46795.34 |
| a,n | -46178.97 |
| v,n | -46741.94 |
| a,v,n | -47170.71 |

**Figure 1:**
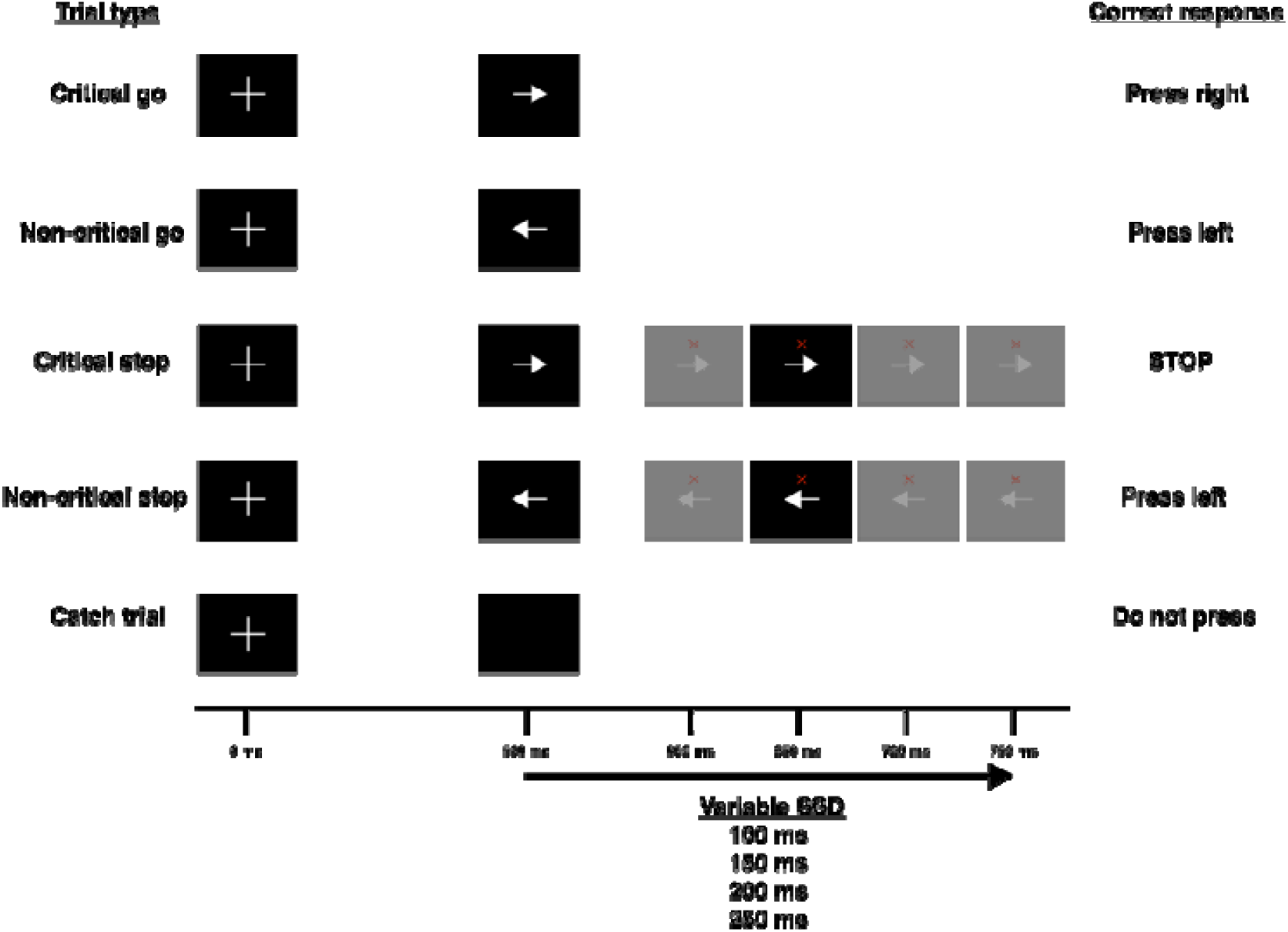
The Conditional stop-signal task (CSST). The different trial types on the CSST with their appropriate responses (critical direction is right). All stimuli stay on the screen until the next stimulus appears. The stop signal delay (SSD) changes between one of four values depending on the performance of the previous critical stop trial.

**Figure 2:**
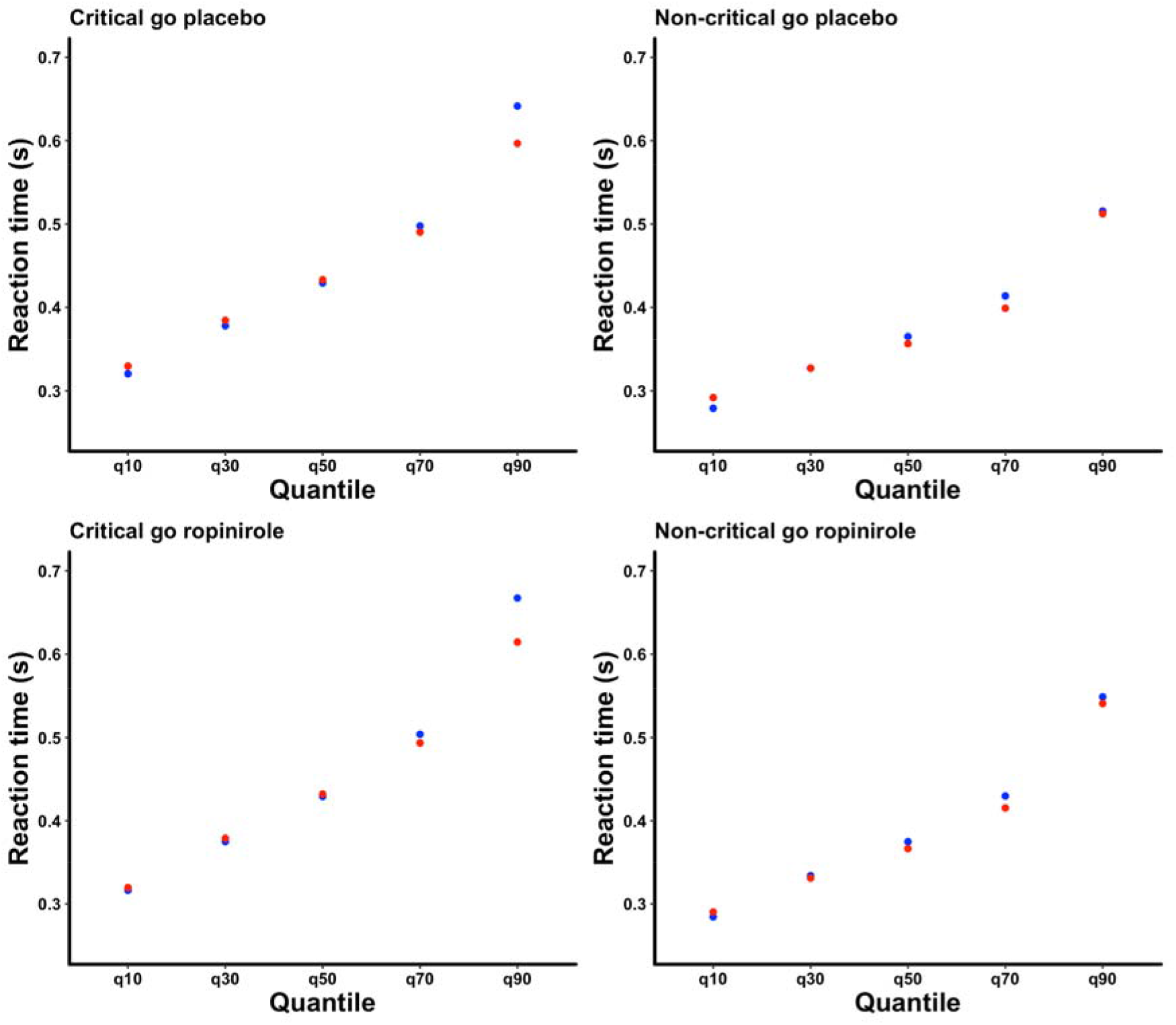
Real and HDDM-predicted reaction times. Plots show predicted (red) and real (blue) reaction times organised by quantile, for each trial/drug permutation. Reaction times are simulated using the HDDM model with the lowest DIC value.

**Figure 3:**
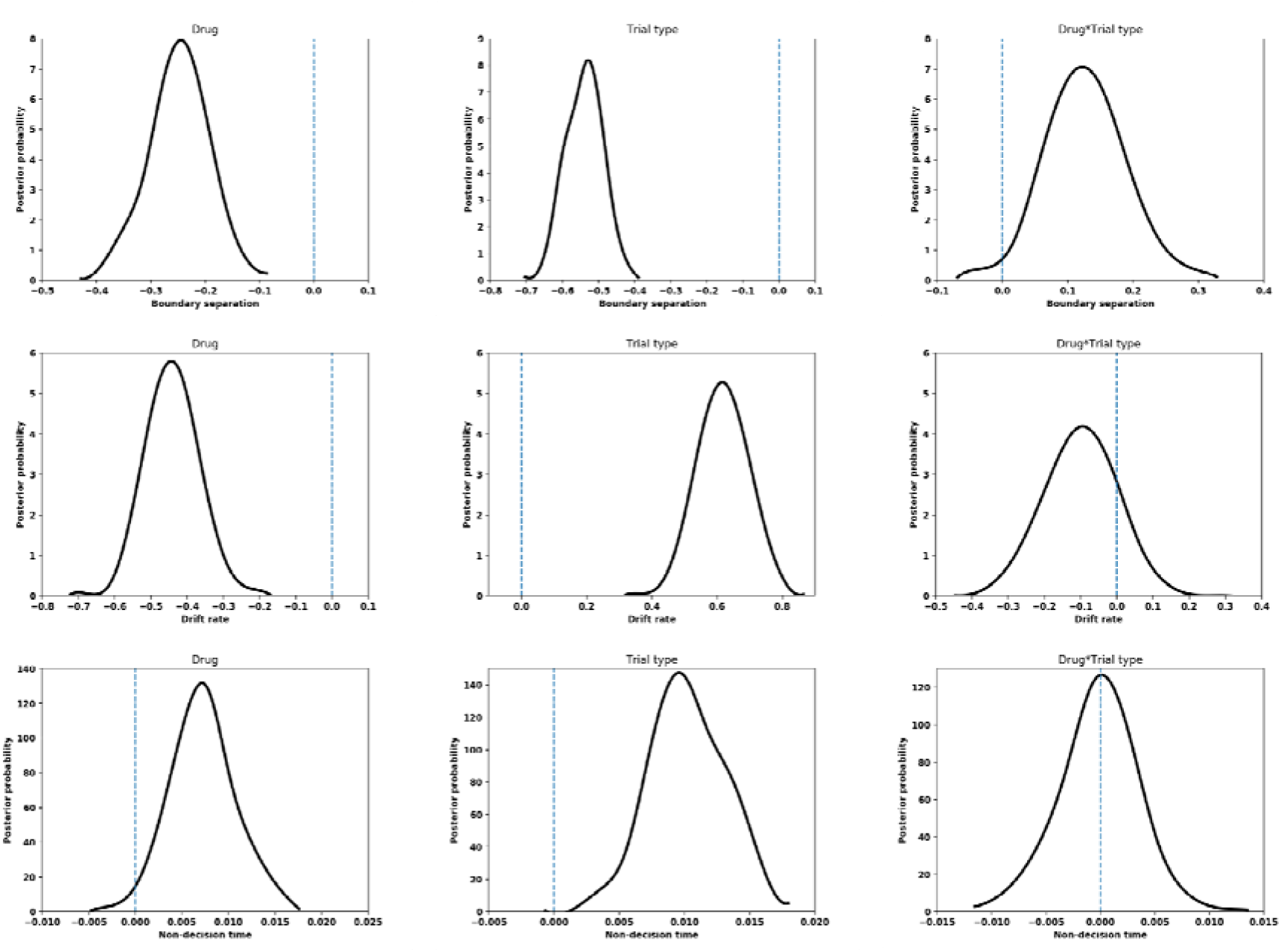
Drift-diffusion model parameters between critical and non-critical Go trials for participants on ropinirole or placebo. Posterior probability distributions are plotted for drug (ropinirole), trial type (non-critical) and an interaction between the two. The blue, vertical line represents the no-effect value. Statistical significance is confirmed if >95% of the distribution lies outside of this line.

From the behavioural analysis, we saw that the RDE was smaller on ropinirole than on placebo; indicating that participants engaged in less proactive inhibition on ropinirole. To investigate how this could arise, we looked for interaction effects between drug and trial type for each DDM parameter. We found that ropinirole specifically impaired the ability to raise the decision threshold on trials when stopping might be required (posterior probability 98.60%). A significant interaction between drug and trial type was not present for drift rate (posterior probability 85.43%) or non-decision time (posterior probability 48.99%).

## Discussion

### A single dose of ropinirole reduces/impairs proactive inhibition in healthy participants

We present data showing how proactive and reactive inhibition are modulated after administration of a single dose of ropinirole, a dopamine agonist with relatively high D_3_ affinity. The data show that proactive inhibition is reduced with administration of ropinirole, indexed by a significant decrease in the RDE. This was accompanied by an increase in the number of go discrimination errors on ropinirole relative to placebo. This is a rather surprising finding since the imperative stimuli are unambiguous, but consistent with a previous report that dopamine replacement in PD resulted in more perceptual decision-making errors on a random motion dots task [28]. By contrast, reactive inhibition as measured by the SSRT, was unaffected by ropinirole administration.

Traditionally, observational and correlative studies have indicated a role of dopamine in response inhibition. Patients with PD (acting as a proxy for chronic dopamine deficiency) show impairments in a variety of tasks probing response inhibition [19], although proactive inhibition seems unaltered in untreated patients [20]. Functional neuroimaging studies show an inverse relationship between dopamine receptor (D_1_ and D_2_) availability and response inhibition [10,29–31]. Interventional studies have shown similar results, with dopaminergic antagonism using haloperidol, resulting in impaired reactive inhibition [32] and dopaminergic agonism using cabergoline or methylphenidate, resulting in improved reactive inhibition [33,34].

A role of ropinirole in response inhibition has been previously explored in healthy adults. In that study, participants were randomised to receive a placebo, 0.5 mg or 1 mg of ropinirole and asked to perform two tasks of response inhibition: a the stop signal task to measure reactive inhibition and a balloon analogue risk task, measuring decisional inhibition, which could be regarded as a form of proactive inhibition. Interestingly, this group also genotyped their participants to derive a dopamine genetic risk score (made up of *DRD1, DRD2, DRD3, DAT* and *COMT* polymorphisms), which indexed basal dopamine neurotransmission. They found that ropinirole modulated both forms of inhibition depending on individuals’ genetic risk score and concluded, as have others [35,36], that dopamine follows an inverted-U shape dose-response relationship with regard to response inhibition [16]. Whilst we did not measure our participants’ dopamine genetic risk score, our results on motor, rather than decisional inhibition reinforce the finding that dopamine agonists modulate proactive response inhibition. It may be the case the underlying genetic susceptibility determines the risk of impulsivity in PD. Indeed, the predictability of ICD generation in PD patients on dopamine replacement is increased when genes associated with ICDs are included in the prediction model [37].

Although the aforementioned studies have informed the role of dopamine in reactive inhibition, the literature concerning dopamine and proactive inhibition remains scarce. D_1_ and D_2_ gene polymorphisms have been shown to predict engagement of proactive inhibition (measured as post-error slowing in a go/no-go task), such that increased D_1_ and decreased D_2_ receptor expression are associated with increased proactive inhibition [38]. Given that ropinirole has a relatively high affinity for D_3_ receptors, our work extends the mediators of proactive inhibition to include D_3_ receptors, as well.

### Ropinirole resulted in less context-specific modulation of boundary separation

We investigated the strategy that participants employed when stopping might be required. Under both ropinirole and placebo, our participants decreased their non-decision time, reduced their drift rate and increased their boundary separation on trials when stopping might be required, the latter confirming previous literature [20]. We then asked what specific effects ropinirole administration had on changes in strategy when stopping might be required. We found that ropinirole resulted in less context-specific adjustment of the boundary separation when stopping was potentially required on critical trials. This finding was in keeping with the behavioural results, where a significantly smaller RDE and a greater number of commission errors were observed. Interestingly, our results mirror those found by Obeso et al. in patients with PD having undergone therapeutic unilateral subthalamotomy. They used the CSST to investigate proactive and reactive inhibition, and found impaired RDE coupled with a failure to engage in context-dependent adjustment of boundary separation during critical trials after right-sided subthalamotomy [20].

Modulation of the dopaminergic system has been shown to alter strategy used during decision-making. Beste and co-workers used methylphenidate, a dopamine/noradrenaline reuptake inhibitor, to study perceptual decision making in healthy individuals using the random motion dots task. They found that administration of methylphenidate significantly increased drift rates compared to placebo and showed that modulation of the dopaminergic system can selectively modulate evidence accumulation [39]. Polymorphisms in dopamine genes have been related to the strategy used during a selective stop signal task [40]. Hence, in addition to acting as a mediator of reactive inhibition, it seems that dopamine has a role in setting the proactive strategy used during specific tasks.

The speed-accuracy tradeoff is a feature of decision-making, whereby decisions that are made faster suffer from decreased accuracy, whereas those that are more accurate are generally slower. The mechanisms by which this tradeoff occurs have been explained using DDMs, with increases in boundary separation resulting in greater accuracy and slower reaction times [41]. Winkel et al. have investigated the role of a dopamine agonist, bromocriptine, on the speed-accuracy tradeoff using the random motion dots task. They failed to find an effect of bromocriptine on boundary separation in mediating the tradeoff [42]. Crucially, the dopamine receptor affinity profile of bromocriptine differs from that of ropinirole; whilst both agents have a high affinity for D_2_ receptors, ropinirole has a relatively higher affinity for D_3_ receptors than bromocriptine. Also, the random motion dots task primarily alters decision-making via changes in drift rate via manipulation of motion coherence of the dots, whereas in the CSST, proactive inhibition is modulated by a combination of strategies. It may be the case that dopamine does indeed have the potential to mediate the speed-accuracy tradeoff via D_3_ receptors. These studies and our own highlight that the role of dopamine in perceptual decision-making is complex, with interactions between neurotransmitter/receptor specificity and design of behavioural tasks.

## Limitations

Although ropinirole has a relatively high affinity for D_3_ receptors, we are aware that it acts on the family of D_2_-like receptors, which include D_2_, D_3_ and D_4_ receptors [43]. Therefore, we are unable to pinpoint the effects of ropinirole on proactive inhibition specifically on D_3_ receptor activity. Whether our findings can be generalized to all dopamine agonists remains to be established; the effect observed in this study may be specific to ropinirole. Future studies could therefore aim to explore a number of dopamine agonists with varying D_3_ receptor activity. If causative, then deficits in proactive inhibition should scale with dopamine agonist D_3_ receptor affinity. Whilst we have drawn parallels between our findings and impulsivity in PD, we note two important caveats; our participants have a preserved dopaminergic state, whereas patients with PD have variable dopaminergic denervation in motor and limbic regions. Furthermore, the doses received by patients with PD typically exceed those used in this study. In view of these differences, conclusions of acute dopaminergic administration in healthy populations cannot be extrapolated to patients with PD. Instead, we encourage specific investigation into the role of dopamine agonists in PD.

## Conclusions

Our results from a placebo controlled, double-blind study provide novel evidence that an acute dose of ropinirole selectively reduces proactive inhibition in healthy participants. The results of the HDDM further indicate that ropinirole results in less context-specific adjustment of the boundary separation when stopping is potentially required. These results may have implications for the potential of this dopamine agonist with high D_3_ affinity to induce impulse control disorders in susceptible patients with Parkinson’s disease.

## Acknowledgements

We would like to thank all participants who gave up their time to take part in this study.

## Author contributions

All authors contributed in the design of the study. All authors were involved in the drafting and revisions of the manuscript. TF provided the ropinirole and placebo used in the study. VR collected and analysed data. All authors have approved this manuscript.

